# Effects of species traits and abiotic factors during the stages of plant invasions

**DOI:** 10.1101/2020.04.20.050278

**Authors:** Daniel Vedder, Ludwig Leidinger, Juliano Sarmento Cabral

## Abstract

1. The success of species invasions depends on multiple factors acting over the four invasion stages transport, colonisation, establishment, and landscape spread. Each of these stages is influenced simultaneously by particular species traits and abiotic factors. While the importance of many of these determinants has already been investigated in relative isolation, they are rarely studied in combination and even then mostly ignore the final phase, i.e., landscape spread.
2. Here we address this shortcoming by exploring the effect of both species traits and abiotic factors on the success of invasions using an individual-based mechanistic model, and relate those factors to the stages of invasion. This approach enables us to explicitly control abiotic factors (temperature as surrogate for productivity, disturbance and propagule pressure) as well as to monitor whole-community trait distributions of environmental adaptation, mass and dispersal abilities. We simulated introductions of plant individuals to an oceanic island to assess which abiotic factors and species traits contribute to invasion success.
3. We found that the most influential factors were higher propagule pressure and a particular set of traits. This invasion trait syndrome was characterized by a relative similarity in functional traits of invasive species to natives, while invasives had on average higher environmental tolerances, higher body mass and increased dispersal abilities, i.e., were more generalist and dispersive.
4. Our results highlight the importance in management practice of reducing the import of alien species, especially from similar habitats.

## Introduction

Species invasions are highly complex phenomena, influenced by several interacting factors, such as species traits, disturbance, or evolutionary history (Theoharides & Dukes, 2007). Gaining an understanding of these factors is necessary to understand the whole invasion process (Fleming & Dibble, 2015) and establish effective countermeasures (Mehta, Haight, Homans, Polasky, & Venette, 2007). Yet, the relative importance of various factors are difficult to derive from studies focussing only on single invasion events (Catford, Jansson, & Nilsson, 2009). Considering the impending global change scenarios and increased rate of biotic exchange, however, generalizable findings about biological invasions are still urgently needed (van Kleunen et al., 2015).

In the last decades, a number of single factors could be identified that contribute to the success of species invasions. A prominent role falls to the number of introduced individuals, i.e., propagule pressure, as it ensures minimal viable population sizes (Lockwood, Cassey, & Blackburn, 2005). Other abiotic factors such as enhanced productivity and increased disturbance have also been suggested to facilitate invasions in some circumstances (Huston, 2004). Beyond these abiotic factors, species traits may also determine invasibility. Arguably the most obvious of these traits is sufficient pre-adaptation to the abiotic environmental conditions of the invaded habitats (Carboni et al., 2016). Furthermore, invasive species need to be able to compete with native species to establish (Hui et al., 2016). Lastly, increased dispersal abilities and broad environmental niche preferences, i.e. generalism, will enable alien species to spread (Rejmánek, 2000). All these invasion factors and stages vary in their level of expression, depending on the system and taxa.

Theoharides and Dukes (2007) put forward a helpful framework integrating many of these factors and stages. They divide the entire invasion process into four stages: transport, colonisation, establishment, and landscape spread. These stages represent a set of community assembly filters that must be overcome before an alien species may be considered “invasive”. The stages do not represent a strict chronology, but rather a set of interlocking and interdependent factors and processes. Hence, for a species to become invasive, it must (a) arrive in sufficient numbers, (b) be adapted enough to the environment to survive, (c) overcome the competition of native species and reproduce, and (d) disperse to establish new neighbouring populations (Theoharides & Dukes, 2007). The main filters involved in these stages are Allee effects, environmental filtering, biotic resistance, and dispersal ability, respectively. Each of these may be influenced by species traits as well as environmental conditions, as illustrated in fig. 1. Concurrent with this framework, it has been demonstrated that the combination of both abiotic factors and species traits has considerable effects on the success of invasions (e.g. Küster, Durka, Kühn, & Klotz, 2010; Mata, Haddad, & Holyoak, 2013; Thuiller, Richardson, Rouget, Proches, & Wilson, 2006). However, many studies in the invasion literature still treat species traits and environmental conditions separately, and frequently only consider one factor at a time. To address this shortcoming, Catford et al. (2009) proposed an experimental design that varies propagule pressure, species composition, and abiotic conditions in a full-factorial setup to assess their relative importance for invasion success. An experimental approach like this will be necessary to arrive at a generalised understanding of the invasion process. However, previous experimental studies focused mainly on the earlier stages of invasion and thus, generalized insights that include landscape spread are still missing (cf. Alzate, Onstein, Etienne, & Bonte, 2020; Kempel, Chrobock, Fischer, Rohr, & van Kleunen, 2013).

**Figure 1:**
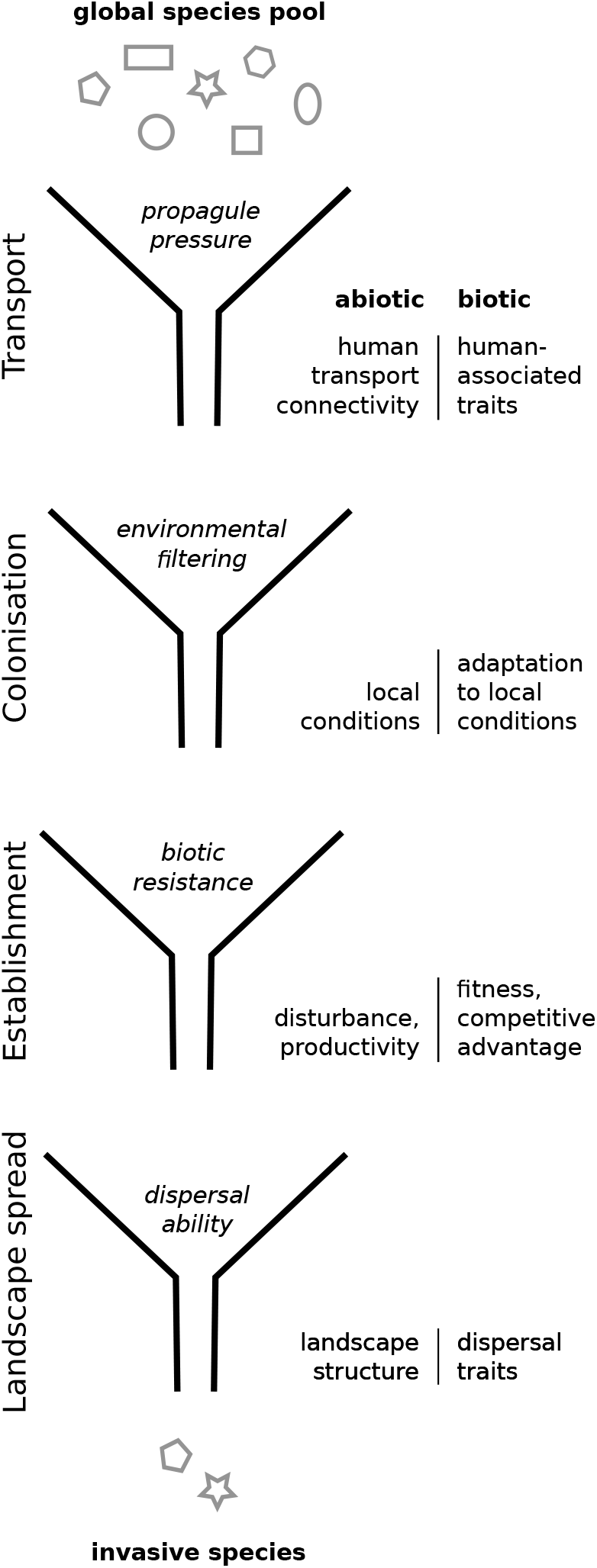
The four stages of invasion (from top to bottom) with their associated filters and abiotic (left label) and trait (right label) factor groups (Theoharides & Dukes, 2007). Apart from the transport factor groups, all factor groups were explicitly modelled (transport factors were combined in a single “introduction” process.)

For generalizing invasion processes, islands can be useful model systems. Firstly, islands are highly susceptible to invasion-related degradation. For example, invasive predators have caused multiple species extinctions on islands (Doherty, Glen, Nimmo, Ritchie, & Dickman, 2016), and there have been observed cases of complete “invasional meltdown” after native keystone species were displaced (O’Dowd, Green, & Lake, 2003). Secondly, their small size, isolation, and comparatively simple ecological dynamics mean that islands are popular study systems in ecology in general (Patiño et al., 2017). These characteristics therefore lend themselves to comprehensively study biological invasions in the context of the stages of invasion (Theoharides & Dukes, 2007). Unfortunately, even for islands, exhaustive data to investigate invasion factors are difficult to obtain and conducting systematic experiments is often unfeasible.

As an alternative, mechanistic models offer a powerful approach to supplement field studies. Such models have previously been used in invasion biology, although usually in the context of specific invaded sites (e.g. Buckley, Briese, & Rees, 2003). However, they also hold great promise for exploring fundamental processes within a more generalised setting (Cabral, Valente, & Hartig, 2017; Grimm & Railsback, 2005; Leidinger & Cabral, 2017), and are very useful for gaining a mechanistic understanding of complex ecological patterns (Grimm & Railsback, 2011). The fact that mechanistic models allow both complete control over all environmental variables, as well as complete knowledge of every individual’s traits makes them an ideal tool to help us better understand the intricacies of the invasion process.

Here, we therefore used a recently developed trait-explicit, individual-based mechanistic model of island plant communities (Leidinger & Cabral, 2020) to investigate the relative roles of propagule pressure, productivity, and disturbance, as well as the traits of invasive species relative to natives. Guided by the framework of Theoharides and Dukes (2007), we wanted to know which factors and trait syndromes increase the success of alien species during the stages of invasions on islands (cf. Pyšek et al., 2015). The specific questions we asked are found in table 1, covering all stages of invasion and including both trait-based and environmental factors. Our experimental setup enabled us to go both broad (covering a wide range of ecological factors) and deep (analysing the resulting species communities down to the trait level) in our investigation of the invasion process. We find that propagule pressure and trait syndromes relative to the native community are the most influential factors promoting invasion success, and discuss how these factors relate to the stages of invasion.

**Table 1:** Questions regarding species invasions to be explored with the model.

| No. | Stage | Question | Expectation |
| --- | --- | --- | --- |
| 1 | *Transport* | How important is propagule pressure for invasibility and why? | Very, because it helps to establish a minimum viable population (Lockwood et al., 2005) |
| 2 | *Colonisation* | Do invasives share a common trait space with native species? | Yes, as they need to be sufficiently adapted to the environment (Carboni et al., 2016) |
| 3 | *Establishment* | How do productivity and disturbance affect invasibility? | Maximum invasibility should be reached at high productivity and high disturbance levels (Huston, 2004) |
| 4 | *Establishment* | Do invasives have a higher fitness than natives? | Yes, as they have to outcompete the natives to establish successfully (Hui et al., 2016) |
| 5 | *Landscape spread* | Are invasives more generalistic and dispersive than natives? | Yes, as this enables them to spread after introduction (Rejmánek, 2000) |

## Methods

### The model

We extended the model of Leidinger and Cabral (2020) to simulate species invasions to plant communities on a virtual oceanic island (fig. 2). The island consisted of a 5 *×* 5 grid depicting a radial elevation (and corresponding temperature) gradient. Additionally, there was a linear (lee-luv) precipitation gradient, which is typical for many oceanic islands. We chose to call this second gradient “precipitation” for simplicity, although it could also be interpreted as any other environmental characteristic like soil type. Each grid cell was assumed to be one hectare in size, with a biomass carrying capacity of two tonnes. Each cell could hold its own community, comprised of individuals belonging to one or more species. Individual body sizes could range between 150 g and 1.2 tonnnes, which puts our system along a range between grassland and shrubland (cf. Deshmukh, 1984). Island geometry and extent were chosen arbitrarily to ensure computational feasibility and to provide different environmental combinations, to increase coexistence of native species (see Armstrong & McGehee, 1980).

**Figure 2:**
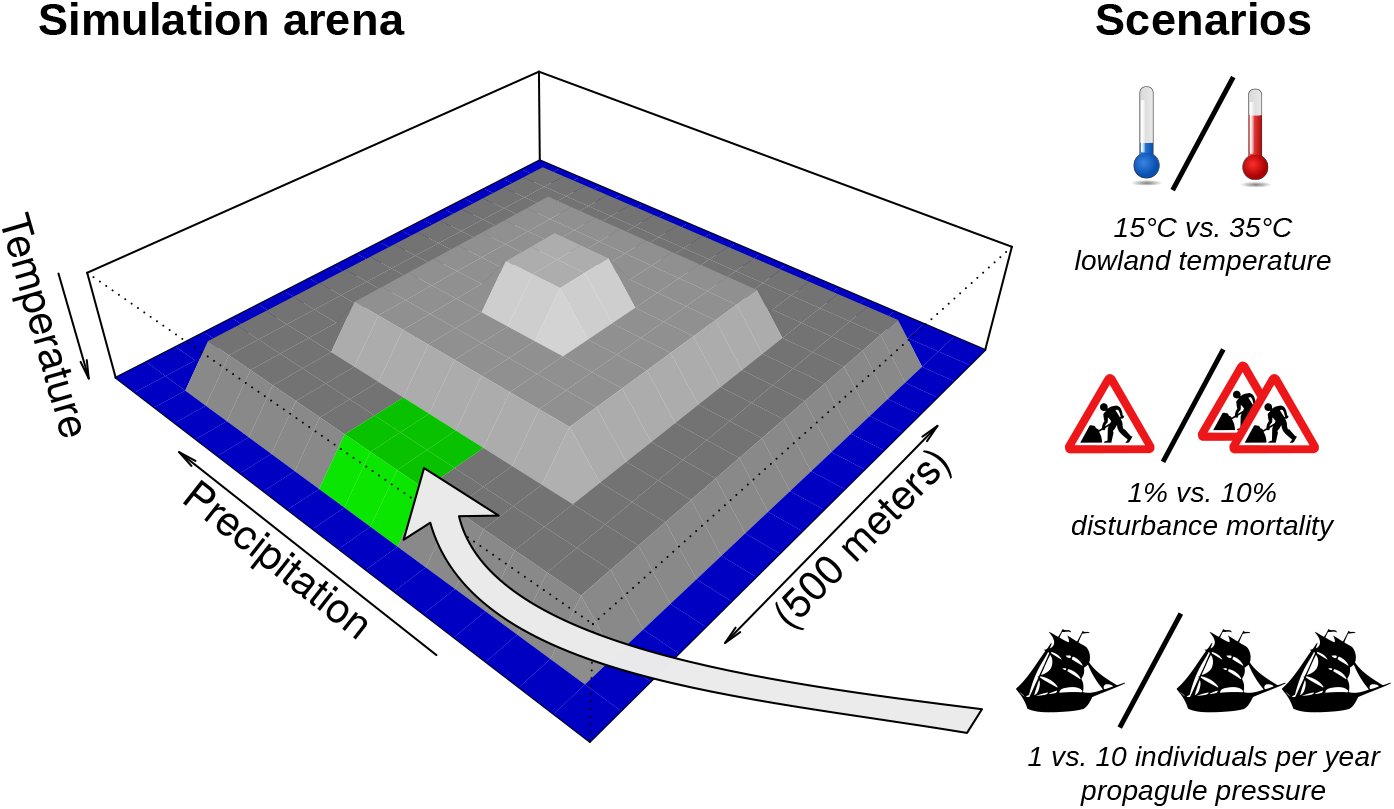
Schematic representation of the simulated island. Temperature decreased with altitude, giving a radial gradient with a step size of 2°C per unit of height (greyscale). A second gradient of an abstract environmental resource (termed “precipitation” for simplicity) was applied longitudinally. The green grid cell denotes the point of entry for alien species. Pictograms show the three factors that were varied in the experimental setup, namely temperature, disturbance, and propagule pressure.

In the model, each individual had a genome consisting of multiple genes that code for a set of traits, which in combination determine the individual’s phenotype. These traits were functional for one or more processes (i.e. traits as parameters of biological functions, table 2). They encompassed environmental optima and tolerances to temperature and precipitation conditions, seed size, reproductive (adult) size, and mean and shape parameters of a logistic dispersal kernel (Bullock et al., 2017). Reproduction only takes place with adult members of the same species within the same grid cell and includes genetic recombination.

**Table 2:**
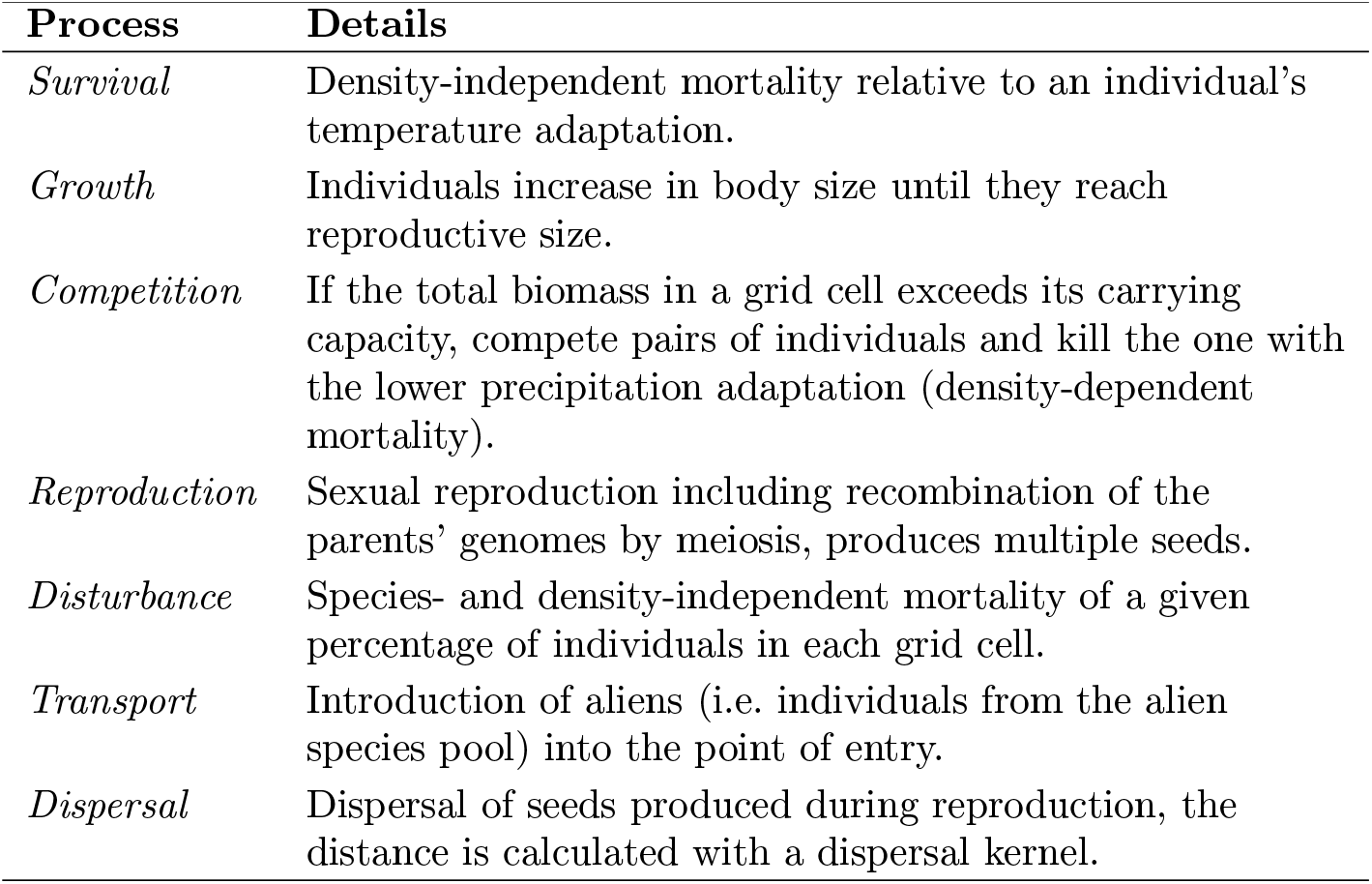
Ecological and life-history processes included in the invasion model, in the order of execution.

| Process | Details |
| --- | --- |
| *Survival* | Density-independent mortality relative to an individual's temperature adaptation. |
| *Growth* | Individuals increase in body size until they reach reproductive size. |
| *Competition* | If the total biomass in a grid cell exceeds its carrying capacity, compete pairs of individuals and kill the one with the lower precipitation adaptation (density-dependent mortality). |
| *Reproduction* | Sexual reproduction including recombination of the parents' genomes by meiosis, produces multiple seeds. |
| *Disturbance* | Species- and density-independent mortality of a given percentage of individuals in each grid cell. |
| *Transport* | Introduction of aliens (i.e. individuals from the alien species pool) into the point of entry. |
| *Dispersal* | Dispersal of seeds produced during reproduction, the distance is calculated with a dispersal kernel. |

The probabilities for growth, density-independent mortality, and seed numbers were determined using the Metabolic Theory of Ecology (MTE, Brown, Gillooly, Allen, Savage, & West, 2004), which links yearly biological rates to body mass and to the local temperature. An individual’s adaptation to the local grid cell temperature additionally scaled the metabolic density-independent mortality probability, whereas adaptation to local precipitation was used to compete individuals when determining density-dependent mortality.

At the initialisation, two species pools were formed with randomly generated species (i.e. each species was characterized by a random combination of genomic and ecological traits). One species pool was used to initialise the island community, and was allowed to stabilise during a “burn-in period” of 500 years. This period length was enough to ensure quasi-equilibrium. Depending on the scenario, native species numbers per island and replicate ranged between one and 15, with a mean of six (Supporting Information, fig. 2). As additions to the original model, the second (alien) species pool had a fixed number of 100 species and was used as a source of alien seeds. After the burn-in period, a fixed number of individuals, regardless of species identity, was drawn from this alien pool and introduced to a specified grid cell on the island (“point of entry”, fig. 2). To further mimic human activities, disturbances also started after the burn-in period, consisting of a given percentage of individuals being randomly removed from each grid cell every year, in addition to the previously mentioned causes of mortality. The model was allowed to run for a total of 1500 years.

For the choice of parameter values, etc., the reader is refered to the full model description in the ODD format (Grimm et al., 2010), found in the Supporting Information. The source code for the model was written in Julia (Bezanson, Edelman, Karpinski, & Shah, 2017), and is available at https://github.com/ lleiding/gemm, along with its documentation.

### Experimental design

We varied the three factors propagule pressure, productivity, and disturbance in a full-factorial design across two levels of each factor to give a total of eight scenarios (cf. Catford et al., 2009; fig. 2). 60 replicates of each scenario were run, resulting in 480 simulations.

We used temperature as a proxy for productivity, varying the base (lowland) temperature between 15°C and 35°C. These values were arbitrary but provided a sufficiently large contrast while maintaining realism. Due to our use of the MTE, higher surrounding temperatures lead to an increase in growth and reproduction rates and thus satisfy the general requirements for productivity (Huston, 2004). Propagule pressure (1 or 10 individuals per year) and disturbance (1% or 10% mortality per year) were explicitly implemented in the model, as described above (cf. Buckley, Bolker, & Rees, 2007; Kempel et al., 2013, for factor values).

### Data recording and analysis

Every year, the model recorded a log file with the mean and variance of each population’s trait values. All data analyses were carried out in R 3.2.3 using `ggplot2` for visualisation (R Core Team, 2017; Wickham, 2016).

To quantify the effect of the varying factors on the success of species invasions (study questions 1 and 3, table 1), we classified species as native, alien, or invasive (species type). Natives were species from the original island species pool that were still extant at the end of the burn-in period. Aliens were all species introduced to the island from the alien species pool. Invasives were the subset of alien species that had established at least one population outside of the point of entry, i.e., had undergone landscape spread (Ricciardi & Cohen, 2007). We then identified all species that became invasive across all replicate runs and summed these up by scenario. During each run, we generated island maps at regular intervals, showing size, location, and species of all populations (e.g. fig. 3).

**Figure 3:**
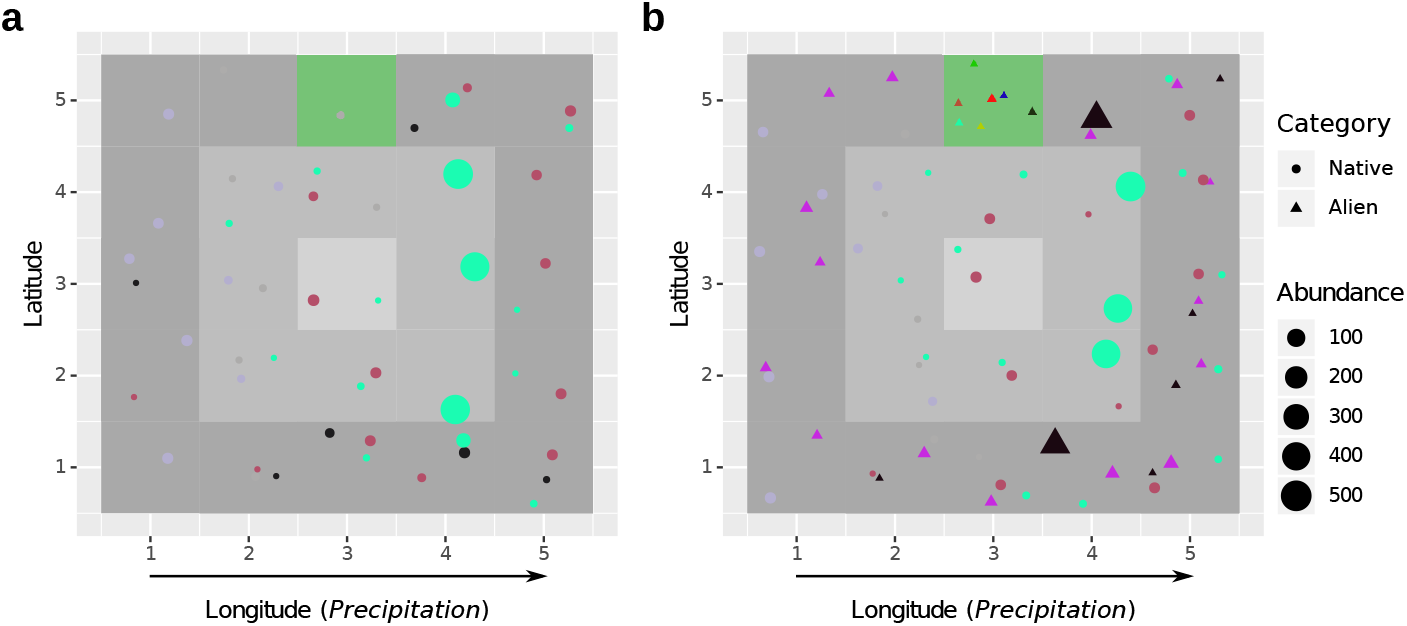
a) Island map after the burn-in period (*t* = 500*a*), and **b)** at the end of the simulation (*t* = 1500*a*) for one example run. Each marker represents one population, colours signify species. Natives are circles, aliens triangles. The green grid cell is the point of entry, grey scale denotes temperature (cf. fig. 2).

To compare the environmental adaptation *A_ind_* of natives, aliens, and invasives, (study question 4, table 1) we combined the model’s internal calculation of adaptation to temperature and precipitation as follows:

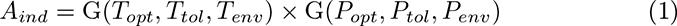

where G(*b, c, x*) is the Gauss function at point *x* with mean *b* and a standard deviation of *c*; *T* is the temperature and *P* the precipitation; *N_opt_* the individual’s niche optimum value, *N_tol_* its niche tolerance, and *N_env_* the actual environmental niche value in the individual’s grid cell.

For trait comparisons between alien, invasive and native species (study questions 2 and 5, table 1), we pooled population data per species type (’alien’, ‘invasive’ or ‘native’) from all scenarios. From this pooled data, we used native populations at the onset of invasion (year 500), and invasive and alien populations over the entire invasion period. The traits we were interested in comprised mean dispersal distance, long distance dispersal, precipitation tolerance, temperature tolerance, adult biomass in grams, and seed biomass in grams. Since precipitation and temperature optima traits were primarily influenced by geography and the particular temperature scenarios, we omitted them from our analysis of the pooled data. Furthermore, we log (*x* + 1)-transformed all trait and adaptation values to improve normality, because the original distributions were left-skewed and contained values *<* 1. We then performed a principal component analysis (PCA) on standardized trait medians of that data to investigate general patterns of trait space by comparing the size and location of 95% confidence interval ellipses corresponding to the different species types. For the PCA, we omitted alien populations, since their increased spread in trait values made comparison of natives versus alien species unfeasible. To assess how the trait characteristics in single traits of native species differed from invasives and identify the most important traits that distinguish native species from invasives, we performed linear mixed models with the particular trait as response, species type as fixed effect, and the specific replicate as random effect, using the R package lme4 (Bates, Mächler, Bolker, & Walker, 2015). We do not report statistical significance of results, since this is mostly meaningless for mechanistic simulation models, as spurious significance emerges by simply increasing replicate number (White, Rassweiler, Samhouri, Stier, & White, 2014).

## Results

A total of 28 species became invasive over all runs, one of them across two different scenarios. We found a strong link between propagule pressure and invasibility, with almost four times as many invasives occurring in high-pressure compared to low-pressure scenarios (fig. 4). Temperature also had a strong influence, with a two- to three-fold difference between levels. There was no clear relationship between invasibility and disturbance.

**Figure 4:**
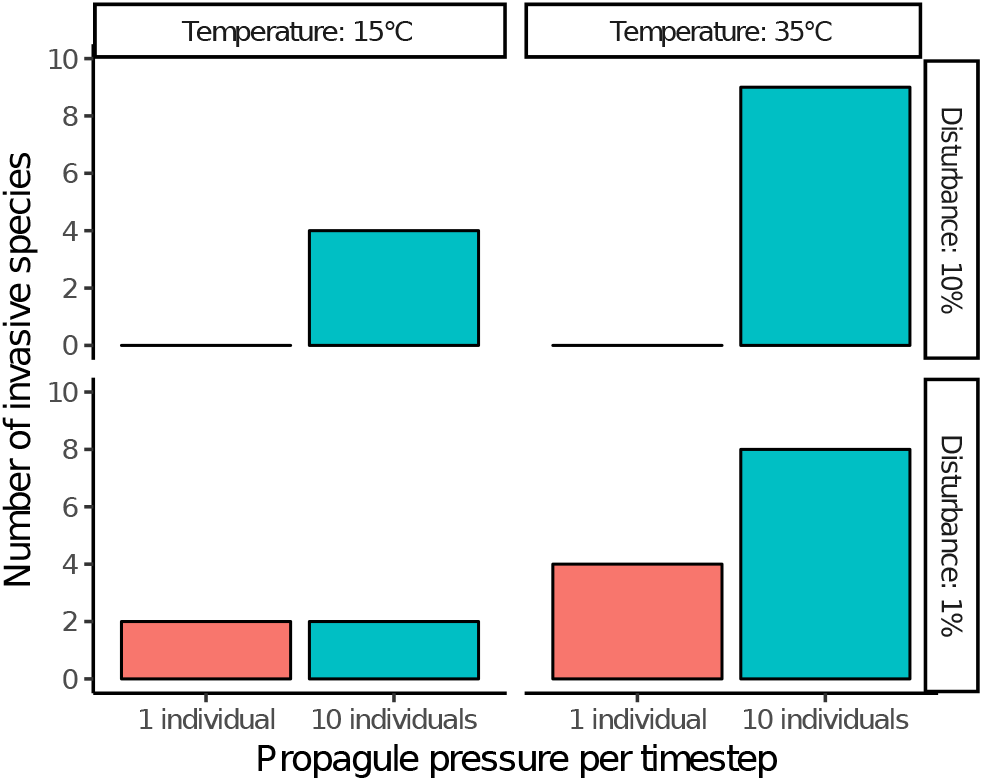
Cumulative number of invasive species observed in each of the eight scenarios (60 replicate runs per scenario).

In terms of the total trait space, invasive populations exhibit a larger spread than natives (fig. 5). The center of the invasive populations’ trait space is shifted along the second PCA dimension towards higher long distance dispersal, higher mean dispersal distance, and higher precipitation tolerance compared to native populations.

**Figure 5:**
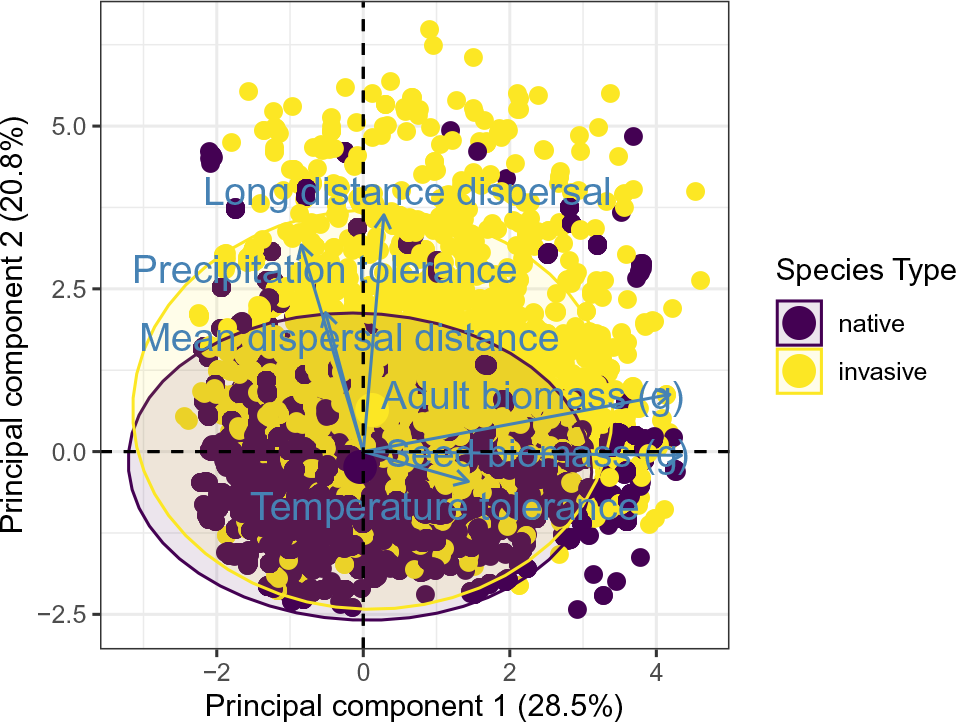
Principle component analysis (PCA) showing the distribution of population trait medians in the trait space. Each axis arrow represents one trait, each marker is one population. All traits were log (*x* + 1) transformed and normalized before PCA calculation. Natives in purple, invasives in yellow. Ellipses represent 95 % confidence envelopes.

The difference in total trait space is associated with specific differences of the particular traits between species category. Specifically, mean dispersal distance (fig. 6a), long distance dispersal (fig. 6b), precipitation tolerance (fig. 6c), temperature tolerance (fig. 6d), and adult biomass (fig. 6e) were all increased in aliens and invasives compared to natives. Seed biomasses were increased in aliens, but slightly decreased for invasives (fig. 6f). For long distance dispersal, precipitation tolerance, temperature tolerance, and adult biomass, the differences in means between natives and invasives were smaller than between natives and aliens. The differences in precipitation and temperature tolerance resulted in the lowest adaptation values for aliens and highest adaptation values for natives, while invasives shows intermediate adaptation value (fig. 6g).

**Figure 6:**
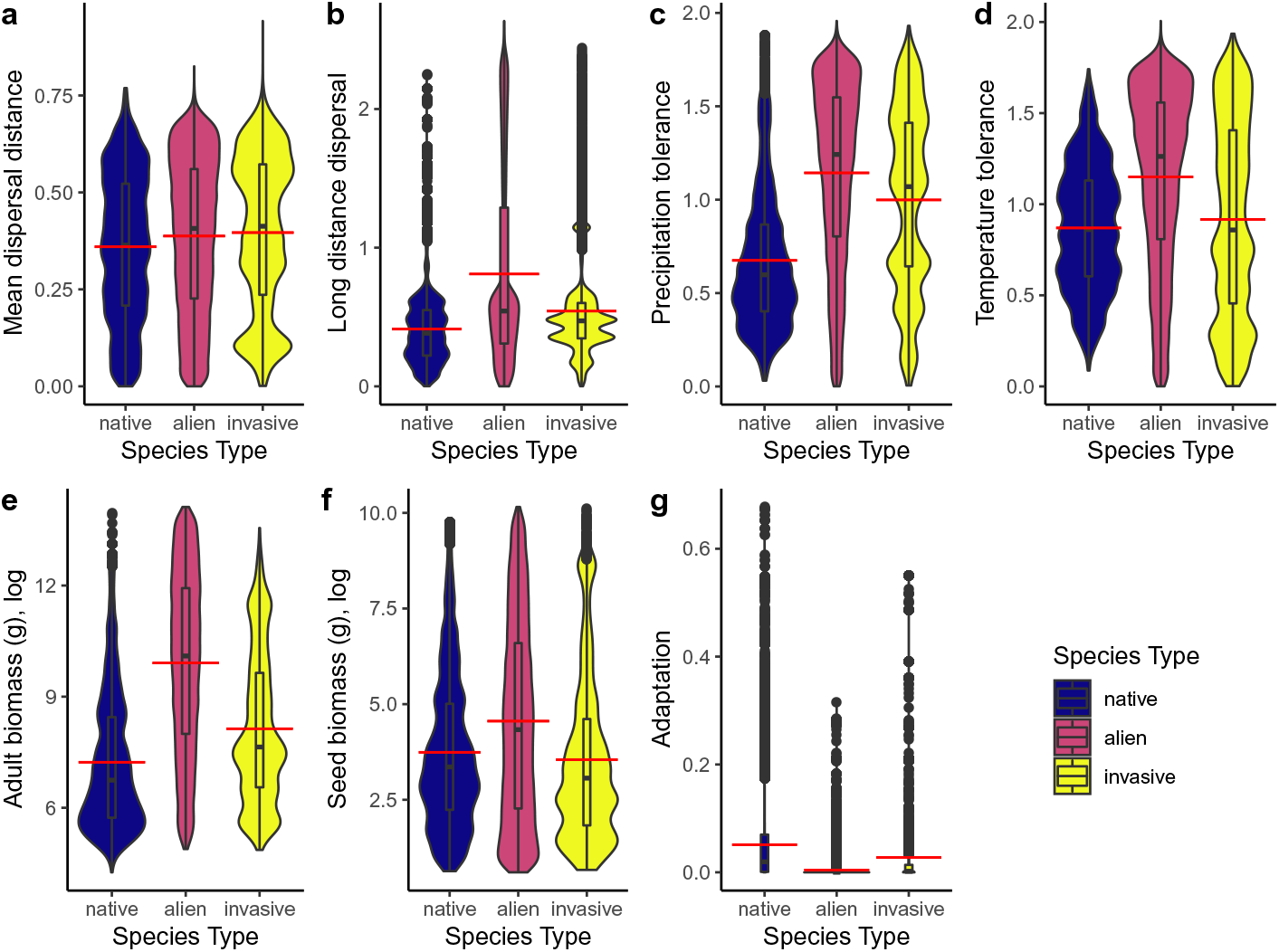
Distributions of single trait values. (a) Mean dispersal distance, (b) long distance dispersal, (c) precipitation tolerance, (d) temperature tolerance, (e) adult biomass in grams, (f) seed biomass grams, (g) adaptation to local temperature and precipitation conditions. All values were log (*x* + 1) transformed before visualisation. Boxes show medians and interquartile range. Red lines highlight the means. Blue: populations of native species, red: populations of alien species, yellow: populations of invasive species.

The results from the linear mixed models revealed what traits were most important to distinguish invasive species from native (table 3). The difference in biomass explained by far the most variance, followed by precipitation tolerance, temperature tolerance, long distance dispersal and seed biomass. Mean dispersal distance explained only a small amount of variance.

**Table 3:**
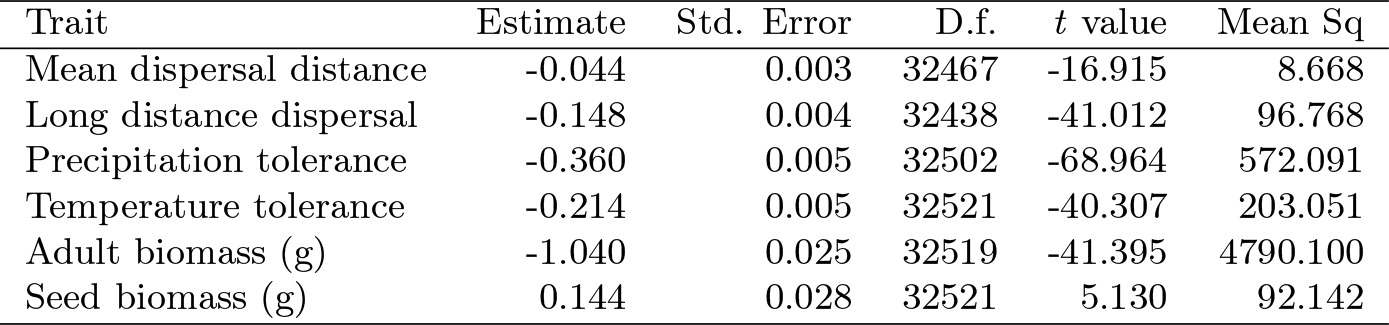
Results of linear mixed-effects model fits comparing the means of species’ traits with the type of species (native or invasive, with invasive as reference) as fixed effect and replicate as random effect. Note that invasive species were more dispersive and generalist with heavier adults, but lighter seeds than native species. Std. Error: standard error, D.f.: degrees of freedom, Mean Sq: mean squares.

| Trait | Estimate | Std. Error | D.f. | *t* value | Mean Sq |
| --- | --- | --- | --- | --- | --- |
| Mean dispersal distance | -0.044 | 0.003 | 32467 | -16.915 | 8.668 |
| Long distance dispersal | -0.148 | 0.004 | 32438 | -41.012 | 96.768 |
| Precipitation tolerance | -0.360 | 0.005 | 32502 | -68.964 | 572.091 |
| Temperature tolerance | -0.214 | 0.005 | 32521 | -40.307 | 203.051 |
| Adult biomass (g) | -1.040 | 0.025 | 32519 | -41.395 | 4790.100 |
| Seed biomass (g) | 0.144 | 0.028 | 32521 | 5.130 | 92.142 |

## Discussion

Although not explicitly imposed by the model, we observe the filtering functions of the stages transport, colonisation, establishment, and landscape spread (fig. 1). Therefore, the general patterns of our model results conform to current knowledge from the empirical and theoretical literature (Theoharides & Dukes, 2007). In line with the requirements of the pattern-oriented modelling paradigm (Grimm & Railsback, 2011), these emergent patterns are found at different ecological levels (individual, population, community, and metacommunity).

### Propagule pressure

Concerning the question on the importance of propagule pressure for the success of invasions (question 1, table 1), our findings mirror ample evidence from empirical studies, including macroecological analyses (e.g. Carr, Hooper, & Dukes, 2019; Seebens et al., 2018). Indeed, propagule pressure is well-known as the leading driver of invasion success in the current literature (Cassey, Delean, Lockwood, Sadowski, & Blackburn, 2018; Lockwood et al., 2005). This is mainly due to Allee effects in introduced populations (Keitt, Lewis, & Holt, 2001; Taylor & Hastings, 2005). Only sufficiently large populations will grow fast enough to overcome adverse abiotic conditions and inter-specific competition by native species. As described in Allee effects theory, this relation of population growth to density is typically hump-shaped, and will decrease again beyond a critical density (Courchamp, Clutton-Brock, & Grenfell, 1999; Stephens, Sutherland, & Freckleton, 1999). Thus, increasing propagule pressure will not increase invasion success indefinitely.

Indeed, Cassey et al. (2018) found a sigmoidal relationship between propagule pressure and establishment success. This was also reflected in additional post-hoc experiments with our model. With propagule pressure increased to 100 individuals per year, we did not observe an increase in the number of successful invasions (Supporting Information, fig. 1). Unfortunately, a direct comparison of our values with those of Cassey et al. (2018) is difficult. This is because Cassey et al. (2018) do not relate their measure of propagule pressure to time, but assume a single initial release, which runs counter to the design of our study. Rather, the repeated introductions in our model increase our *de facto* propagule pressure beyond the one or ten individuals introduced in one year; but the stochastic nature of introductions makes it impossible to quantify exact values per species. Nevertheless, the results of our subsequent simulations indicate that we did reach propagule saturation as well. The reason for this is likely increased intra- and interspecific competition among juvenile alien individuals in already saturated communities. This saturation is an effect of our initial simulation conditions, where we initialize communities with more species populations than they will eventually hold. Given that real islands are known to have lower species richness than continents (Whittaker & Fernández-Palacios, 2007) and are thus likely un-saturated in terms of species numbers, they will therefore be much more susceptible to increased introductions than saturated continental systems. This highlights again the vulnerability of island biota to global change processes.

### Temperature and disturbance

The interaction between temperature (our surrogate for productivity) and disturbance was not as straight-forward as we had expected (question 3, table 1). The dynamic equilibrium model (Huston, 2004) predicts that invasibility approximately increases with native diversity, which is high when both productivity and disturbance are high, or when both are low. This is because high productivity with low disturbance leads to population extinction through competitive exclusion, while low productivity with high disturbance means small populations that go extinct stochastically (Huston, 2004). Indeed, as described by Huston (2004), we did observe a clear peak of mean native species rich-ness in the low-temperature, low-disturbance scenarios (Supporting Information, fig. 2). Despite this, maximum invasibility did not coincide with either low-temperature/low-disturbance or high-temperature/high-disturbance scenarios, but was driven almost entirely by temperature (fig. 4). This apparent mismatch of our model with theory may raise some questions at first, but can be explained by a closer look at disturbance and its influence on invasion processes.

The interplay between productivity and disturbance and its effect on species richness and composition has long been discussed in the theoretical literature (e.g. Catford et al., 2012; Chesson, 2000) and shown in multiple empirical studies (e.g. Huebner, Regula, & McGill, 2018; O’Connor, Falk, Lynch, Swetnam, & Wilcox, 2017). However, these interactions are subject to several preconditions. Firstly, Buckley et al. (2007) point out that disturbance may increase the invasibility of a habitat, but can also be a cause of mortality for alien species, reducing invasibility again. Thus, if disturbance affects natives and aliens equally, these two contrary effects may cancel each other out, leaving no net change. This was the case in our model, as disturbance-driven mortality was species-agnostic. Secondly, the effects of disturbance on native and alien communities are likely to change over the full gradient of disturbance intensity (Catford et al., 2012). Analysing that many disturbance levels was beyond the scope of this study, however. Therefore, our experiment only reflects two points on this gradient and may thus give an incomplete picture. And thirdly, an invaded community’s response to disturbance can be strongly modulated by the trait composition of its native and alien species (Kempel et al., 2013; Mata et al., 2013). This includes traits that are represented in our model, which will be discussed in the next section.

### Traits of invasive species

Although sometimes questioned in studies on species invasions (cf. Catford et al., 2009, but see Thuiller et al., 2006), our results provide some insights into an “invasion syndrome” relative to the native community and the stages of invasion (question 2, table 1; Novoa et al., 2020). In fact, invasive species are more similar to native species than alien species in terms of their trait characteristics (cf. Küster et al., 2010). This suggests that invasive species have to pass similar environmental filters to natives in order to complete the colonization and establishment stages.

However, this trait similarity is not sufficient to surpass native species in terms of environmental adaptation (question 4, table 1). Instead, invasive species have on average lower environmental adaptation. The fact that some invasive species are still successful, and even end up replacing native species, again highlights the importance of propagule pressure. This might offset some of the maladaptation and the priority effect advantage enjoyed by native species (Chase, 2003). Such interaction of traits and propagule pressure is in line with recent evidence from field and laboratory experiments (Alzate et al., 2020; Kempel et al., 2013). Still, propagule pressure alone does not explain why invasives also manage the fourth phase, landscape spread.

In order for invasives to spread, we expected increased dispersal abilities and a rather generalistic niche strategy (question 5, table 1). Indeed, invasive species in our simulations not only feature increased mean and long distance dispersal, but also have higher environmental tolerances on average (see Grotkopp & Rejmánek, 2007; Schultz & Dibble, 2012). These latter trait characteristics suggest the generalist nature of invasive species, at least in comparison to natives. A larger biomass and thus reduced mortality in invasives additionally makes them stronger competitors (cf. Kempel et al., 2013). This circumstance will also help offsetting relative maladaptation, besides propagule pressure.

Furthermore, the difference in trait characteristics between species categories might also indicate different evolutionary backgrounds. In this respect, our island communities are the result of several hundreds of years of ecological interactions between species and the environment. This is especially evidenced by their decreased dispersal ability, which is a typical island adaptation (Burns, 2019). They are, therefore, sensitive toward invasive species that (1) were not restricted by those interactions, and (2) exhibit trait syndromes different from those represented in the community. Additionally, invasive species are bigger than natives, but not as big as (unsuccessful) alien species. Thus, the invasion syndrome features adaptation to similar conditions as the invaded native community, but sufficiently different other key traits (e.g. biomass), unbounded by the local evolutionary history, to fill areas of the trait space with the least overlap with native species. If this overlap is enough to induce competitive exclusion, superior competitive abilities allow invasive species to outperform, and thus replace, native species, which we could observe in some of our simulations (cf. Flory & Clay, 2010; Pyšek et al., 2012).

### Future considerations

In our simulations, we could not account for most biological interactions except for competition. Mutualisms between native species will likely lead to secondary extinctions, or even extinction cascades, if keystone species get lost (Christian, 2001; Schachtschneider & February, 2013). Trophic interactions can also allow effects like enemy-release of alien species. Therefore, the effects of invasions on our simulated communities might be very conservative and underestimated. We anticipate that the consideration of additional interactions into our modeling framework will further increase the impact of species invasions on the native communities.

In this study, we corroborated the importance of propagule pressure for the invasion success. It is important to point out, however, that previous studies frequently conflated this with the related but separate concept of colonization pressure, i.e., the number of species (rather than individuals) introduced per time (Lockwood, Cassey, & Blackburn, 2009). Although the two may interact and are difficult to differentiate in the field, they do in fact address two quite dissimilar mechanisms: propagule pressure determines how quickly a minimum viable population can be established, whereas colonization pressure increases the chances of introducing a suitable species. Our experiment does not completely disentangle these two aspects, but by using an alien species pool of constant size, we still arguably test the more specific understanding of “propagule pressure”. However, future investigations could use a model such as ours to fully separate the effects of both these factors in designated experiments.

Increasing the runtime of the model may lead to additional insights. A number of studies have pointed to the impact of longer time scales on the invasion process. For example, Pyšek et al. (2015) and Carr et al. (2019) show a link between residence time or sustained propagule pressure and establishment rates. On the other hand, Sheppard and Schurr (2019) observed an increase in biotic resistance over time, which decreases the performance of alien species. Investigating such long-term and large-scale effects of invasion factors further is possible with our model, although this will require significant computational resources.

Although beyond of the scope of this study, the current model allows us to consider genomic traits in the characterisation of species. In a previous study, Leidinger and Cabral (2020) show how environmental variation interacted with the number of genes and genomic variation of communities. Similarly, invasive species might display particular genomic profiles which enable them to quickly adapt to novel environments, for example by high standing variation or phenotypic plasticity (Zenni, Lamy, Lamarque, & Porté, 2014). To increase computational feasibility and preclude possible confounding effects, we chose not to include these effects in the present study. However, we expect future experimental designs to account for variation in genomic traits and to allow for mutations when investigating species invasions.

This study demonstrates the utility of individual-based mechanistic models for understanding biological invasions. Our results hold relevance for policy and management, as they reinforce the importance of reducing the import of alien species. This is true of alien species in general, but even more so for those showing high environmental tolerances and dispersal abilities, as well as those coming from habitats with similar conditions as native ecosystems.

## Data Accessibility

All data should be reproducible by using the simulation codes found at https:// github.com/lleiding/gemm.

## Author contributions

DV, LL, and JSC conceived the ideas and designed methodology; DV performed simulations; DV and LL analysed the data; DV led the writing of the manuscript. All authors contributed critically to the drafts and gave final approval for publication.

