## Supporting information for "Effects of species traits and abiotic factors during the stages of plant invasions"

#### 1 Figures

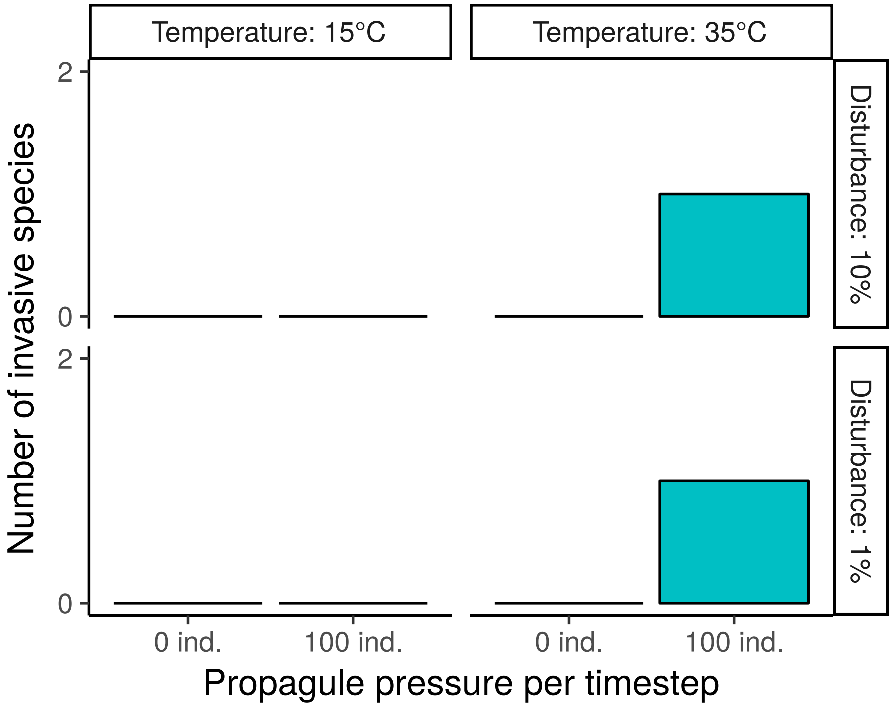


Figure 1: Number of successful invasions per scenario at higher propagule pressure levels (post-hoc simulation data, 10 runs per scenario).

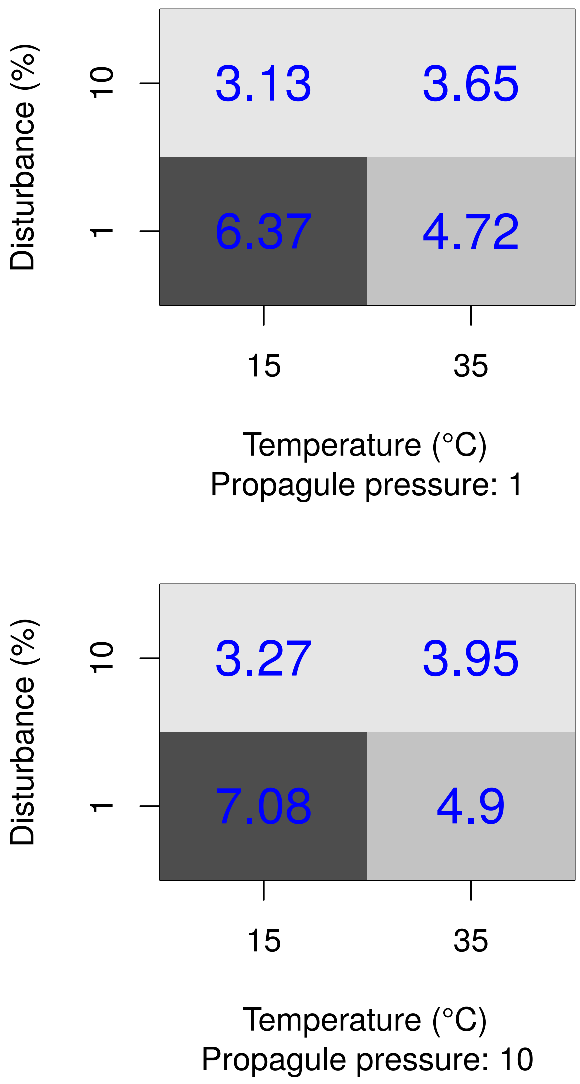


Figure 2: Average native species richness per scenario (main simulation data, 60 runs per scenario).

### 2 Model description

The model description follows the ODD (Overview, Design concepts, Details) protocol (Grimm et al., 2006, 2010).

#### 2.1 Purpose

This model is designed to simulate a (meta-)community of plant-like individuals. For this, the model considers factors and processes across genetic, population and ecological levels. The model is able to produce several patterns across genetic, individual, population and (meta-)community levels, including adaptation and speciation through divergence of populations. Thus, the model expands from basic principles to richer representation of real-world scenarios.

#### 2.2 Entities, state variables and scales

Individuals are the basic entity in the model. Given their attributes and life-history, these individuals most closely resemble plants. Individuals belong to different species, which are characterized by similar ecological traits and identical genetic architecture. The genetic architecture of each individual is comprised of a diploid set of one or more linkage units, which, in turn, combine a set of genes. Linkage units are always inherited in their entirety during the recombination phase of a reproduction event. The higher the number of genes per linkage unit, the higher the degree of genetic linkage. Some of the genes code for one ore more traits (pleiotropy), while a trait can be dependent on more than one gene (polygene). The realized trait value is the mean of all the trait alleles (quantitative trait loci). Traits thus controlled encompass the initial body mass (size) of offspring, $M_s$, the body mass determining onset of maturity and thus reproductive capability, $M_r$, mean dispersal distance, $\mu$, the shape of the dispersal kernel, controlling long-distance-dispersal, $s$, and values representing the optimum and the tolerance (standard deviation) of a physical niche parameter, such as temperature and precipitation ($\overline{T}$ and $\sigma_T$ or $\overline{P}$ and $\sigma_P$, resp.). Alternatively to be controlled by mutable genes, traits can also be fixed. Additionally, individuals carry attributes which describe their bodymass, $M$, and their adaptation to the abiotic environmental conditions (fitness), $F_T$ and $F_P$. Furthermore, every individual carries a Boolean marker used to store whether a given individual has newly arrived to a grid cell or discriminate individuals from the rest of the community.

The base rates for processes governed by the metabolic theory of ecology (Brown et al., 2004) - growth, reproduction, mortality - are global constants. Mutation rate is also a global constant.

Every individual is placed inside an arena of grid cells, each of which has a unique location (coordinates) and is characterized by physical properties such as temperature, precipiation and size (carrying capacity). Over the course of the simulation these properties (location or physical parameters) might change, reflecting geomorphological dynamics. All individuals within one grid cell constitute a community. The characteristics of the grid cells combined with the state of inhabiting individuals constitute the state variables of the model. Additional patterns or summary statistics may be calculated based on these individual information.

Processes and updates are repeated every timestep, while each timestep can be considered as one year.

#### 2.3 Process overview and scheduling

In each discrete, yearly timestep each individual in each grid cell will (in no particular order unless otherwise stated) undergoes the following processes: (1) establishment, (2) density independent mortality influenced by adaptation to temperature, (3) disturbance, (4) growth, (5) competition (individuals are sorted according to their adaptation to precipitation), (6) reproduction (7) mutation of offspring, (8) filtering of unviable individuals, (9) seed dispersal.

Before seed dispersal, alien individuals might be introduced to designated grid cells during an invasion process. After seed dispersal, the physical environment of a grid cell might change. If that happens, all individuals within that cell are marked to undergo establishment again.

Updates to individuals and thus the local communities happen instantaneously after a specific process has been executed (asynchronous updating).

#### 2.4 Design concepts

##### Basic principles

Metabolic theory of ecology (at the submodel level). Adaptive and non-adaptive radiation/evolution (mostly submodel level, but geomorphological change at the system level). Sexual reproduction. Niche theory (both at system and submodel level). Each individual carries unique ecological and/or functional traits, as well as preferences for their physical environments. Resource/energy limitation (carrying capacity: system level property, but invoked at submodel level).

##### Emergence

Species/Populations/ecotypes. Only constrained via genetic properties. Community trait composition. Interplay of physical properties (environment, geographical properties) and within community (competition strength via reproduction, growth, etc.). Species numbers, endemics, speciation rate.

##### Adaption & Objectives

See entities. Traits follow evolution: a trait changes its value randomly within a given phylogenetic constraint. The success of the change (fitness) emerges as the result of adaption to the physical environment and the reproductive success of an individual over its competitors.

###### Learning

N/A

###### Prediction

N/A

###### Sensing

Individuals are directly affected by the properties of their physical environment (e.g., temperature).

###### Interaction

Individuals directly interact when sexually reproducing. However, they are not affected themself by this interaction. Instead, the interaction aims solely at determining the genotype of their offspring. Additionally, competition for resource/energy/space between individuals represents indirect interaction.

###### Stochasticity

Most of the submodels are carried out by all elegible individuals. Some submodels (Survival, Competition, Mutation and Dispersal), however, happen with particular probabilities. In these cases, execution of submodels is decided at random, taking into account individual characteristics, such as body size, fitness, genome size, or dispersal abilities. All decisions inside all of the submodels are stochastic (e.g., number of offspring) to maintain variability and relax assumptions.

###### Collectives

N/A

###### Observation

At the start and end of the simulation and at definable regular time intervals, the properties of all individuals (including the properties of their locations) are recorded and written to files.

#### 2.5 Initialisation

The initialisation step creates lineages with randomly chosen genetic and ecological trait values in each grid cell that is designated to receive an initial community. This encompasses choosing the number of genes for a lineage, the number of linkage units and the within genome variance of trait values. Trait with thus distributed trait values are the distributed randomly among the genes. Population (number of individuals) size of a lineage is determined by the adult body size of individuals from a lineage. At this point all individuals of a population are identical. Values for ecological traits are then varied in each gene where a given trait is found, for all individuals of a lineage. The variation is Normal distributed with the lineage trait value as mean and the product of $\sigma_l$ (phylogenetic constraint) and the lineage trait value as standard deviation. This ensures initial genetic variation within a lineage population. Thus created populations are added to a grid cell's community until the additional mass of another population would exceed the grid cell's carrying capacity. In the experiment configuration file, it is possible to specify other methods of initializing communities, e.g., "single", where each grid cell receives only one species. Whether a grid cell receives an initial community depends on the map definition. At the end of initialisation each of the thus populated grid cells holds one or more different populations, each from a separate lineage.

#### 2.6 Input

At the start of a simulation user defined parameters are read, containing also a definition of the simulation arena (map definition). This definition is provided in a separate plain text file. Within the text file a line at the top containing a single number defines the number of timesteps the arena definition is valid for. Every other non-empty line defines one grid cell with a unique identifier (a number), and the location of the grid cell as two coordinates. Optionally, one can define the type of the grid cell (island or continent), its temperature and precipitation values, and size.

Other optional parameters can be set in a separate configuration file and pertain to defining simulation scenarios:

| Name/Function | Default value | Description |
|---|---|---|
| "avgnoloci" | 1 | average number of loci/copies per gene |
| "biggenelength" | 200, | Sequence length of long genes |
| "burn-in" | 500, | timesteps before invasion starts |
| "cellsize" | 2e6, | maximum biomass per hectare in gramm (cf. Deshmukh, 1984; Clark et al., 2001) |
| "config" | "simulation.conf", | configuration file name |
| "debug" | false, | write out debug statements |
| "dest" | string(Dates.today()), | output folder name |
| "disturbance" | 0, | percentage of individuals killed per update per cell |
| "fasta" | false, | record fasta data? |
| "fertility" | exp(28.0), | global base reproduction rate 23.8 from Brown et al. (2004), alternatively 25.0, default 30.0 |
| "fixtol" | true, | Reproductive tolerance does not evolve |

| Name/Function | Default value | Description |
|---|---|---|
| "global-species-pool" | 100 | size of the global species pool (invasion source) |
| "growthrate" | exp(25.2), | global base growth/biomass production from Brown et al. (2004) |
| "indsize" | "seed", | initialize organisms as seed, adult or mixed |
| "lineages" | false, | record lineage and diversity data? |
| "linkage" | "random", | gene linkage type (random/full/none) |
| "logging" | false, | write output to logfile |
| "maps" | "", | comma-separated list of map files |
| "maxdispmean" | 10, | maximum mean dispersal distance |
| "maxrepsize" | 14, | maximal repsize in grams calculated as exp(maxrepsize) → 1.2 t |
| "maxseedsize" | 10, | maximal seedsize in grams calculated as exp(maxseedsize) → 22 kg |
| "maxtemp" | 313, | max optimum temp in K |
| "minrepsize" | 5, | minimal repsize in grams calculated as exp(minrepsize) → 150 g |
| "minseedsize" | 0, | minimal seedsize in grams calculated as exp(minseedsize) → 1 g |
| "mintemp" | 283, | min optimum temp in K |
| "mortality" | exp(22), | global base mortality from Brown et al. (2004) is 26.3, but competition and dispersal introduce add. mort. |
| "mutate" | true, | mutations occur |
| "mutationrate" | 3.6e10, | one mutation per generation/individual, corrected for metabolic function |
| "nniches" | 2, | number of environmental niches (max. 3) |
| "outfreq" | 100, | output frequency |

| Name/Function | Default value | Description |
|---|---|---|
| “phylconstr” | 0.1, | phylogenetic constraint during mutation and inter-loci variation. scales trait value as sd. |
| “phylo” | false, | record phylogeny? |
| “popsize” | “metabolic”, | initialisation algorithm: metabolic/bodysize/minimal/single |
| “precrange” | 10, | range from 0 for precipitation optimum |
| “propagule-pressure” | 0, | number of non-native individuals introduced per invasion event |
| “quiet” | false, | don’t write output to screen |
| “sdtemp” | 0.0, | SD of temperature change per time step |
| “seed” | 0, | for the RNG, seed $= 0 \rightarrow$ random seed |
| “smallgenelength” | 20, | Sequence length of regular/small genes |
| “static” | true, | mainland sites don’t undergo eco-evolutionary |
| “tolerance” | 0.8, | sequence similarity threshold for reproduction |
| “traitnames” | [“compat”, “dispmean”, “dispshape”, “precopt”, “prectol”, “repsize”, “reptol”, “seedsize”, “tempopt”, “temptol”], | minimal required traitnames |
| “usebiggenes” | true | Whether to use longer genes |

If a parameter value is not specified by the user, the default value for that parameter set in the simulation code is assumed. Global parameter values were either adapted from the literature or found via trying out a range of values to identify combinations that lead to high species coexistence.

#### 2.7 Submodels

##### Establishment

Whenever an individual is new to a grid cell (by recent birth, dispersal event or environmental change), their physical niche preferences are compared with the actual niche properties, e.g., the temperature, $T$, of the present grid cell. The individual adaptation parameter, $A$, is set according to the deviation from the optimum value considering the niche breadth as standard deviation of a

Gaussian curve, i.e., an individual's fundamental environmental niche.

$$A = a \times \exp\left(-(T - \overline{T})^2/(2\sigma_T^2)\right) \tag{1}$$

where

$$a = 1/(\sigma_T \sqrt{2\pi}) \tag{2}$$

##### Competition

If the sum of the community's bodymass exceed the available space, this will pick two individuals at random and remove the one that has lower adaptation to local precipitation, $A_P$. Once total bodymass is below carrying capacity, the procedure terminates.

##### Growth

Given an individual has undergone establishment, an individual changes its size $(M + \delta_M)$ following the metabolic theory and the global base growth rate, $b_0$:

$$\delta_M = b_0 \times M^{\frac{3}{4}} \times \exp\left(\frac{-E_A}{k_B \times T}\right) \tag{3}$$

with $E_A$ as activation energy and $k_B$ the Boltzmann constant. In case this change results in zero or negative body mass, the individual is removed from the community.

##### Density independent mortality / Survival

An individual is removed from the local community with a probability $p_mort$ depending on its size $M$, its adaptation to temperature, $A_T$, and a global base mortality rate $b_mort$:

$$p_mort = \left(1 - \exp\left(b_{mort} \times M^{-\frac{1}{4}} \times \exp\left(\frac{-E_A}{k_B \times T}\right)\right)\right) \times {A_T}^{-1} \tag{4}$$

##### Reproduction and mutation

All individuals that have grown to or beyond their individual reproduction sizes may reproduce. The number of offspring is randomly drawn following a Poisson distribution with mean $N$ determined by the individual's size $M$ and a global base offspring number $N_0$:

$$N = N_0 \times M^{-\frac{1}{4}} \times \exp\left(\frac{-E_A}{k_B \times T}\right) \tag{5}$$

The number of offspring is then multiplied by the seed mass encoded in the parent's genome and this total biomass subtracted from the parental biomass. If the remaining biomass would be equal to or less than 0, the individual will not reproduce. Otherwise, possible mates are selected within the same grid cell based on whether they belong to the same lineage, have reached maturity (which includes having established on the grid cell) and whether their compatibility sequences are sufficiently similar. If a suitable partner is found, both partners produce gametes, i.e., complete haploid sets of all linkage units, where each linkage unit is randomly picked either from the maternal or paternal set. The two gametes, one from each mating partner, comprise the genome for the offspring. At this point, mutations in the offspring's basecode may happen with a set probability $P_m$. In the case of mutation all traits associated with the respective gene will randomly change value $V$ by $\epsilon$, which is normally distributed and has as standard deviation the product of $\sigma_l$, i.e., the phylogenetic constraint, and $V$.

The new individuals' trait values are then calculated as the means of all alleles and the individuals added to the community, with their size set to the initial biomass $M_s$ (seed biomass).

##### Dispersal

After reproduction and mutation, each offspring individual may disperse. For each of these, a new location (i.e. $x$ and $y$ coordinates) is drawn randomly following a logistic distribution with mean and shape parameters (which controls long-distance-dispersal) taken from the individual's traits. If a suitable grid cell is found at the drawn coordinates, the dispersing individual will be placed there and removed from the original community. The removal happens even when there is no destination grid cell to be found. Special attention is paid when the destination grid cell is of island type, while the origin is on the mainland and the simulation runs in static mode. In this case the dispersing individual is copied to the new destination instead of moved.

##### Habitat change

If enabled, both environmental habitat parameters - temperature and precipitation - change values throughout the simulation arena. The amount and direction of change is the same for all grid cells across the landscape. Changes to temperature and precipiation happen independently from one another. The change is randomly drawn from a Normal distribution with the current value as the mean and a user defined standard deviation.

##### Disturbance

If enabled, a set percentage (determined by the `disturbance` setting) of individuals in each cell is killed each turn, regardless of species or individual properties. Disturbance only begins after the `burn-in` period.

##### Invasion

If enabled, a global species pool is initialised with `global-species-pool` random species. After the `burn-in` period, `propagule-pressure` individuals from this pool are randomly selected and copied to the simulation arena's point of entry (multiple selection of the same species per turn is possible).

#### 2.8 Output/Calculation

The main simulation data output is stored in two separate formats. The first is a table containg data characterising the individuals. Each line represents on individual. The columns describe an individual's current state. This is characterised by location, environmental conditions, ecological traits and summary of the genetic architecture. Additionally or alternatively to the individual level data, the data can be summarized at the population level (i.e. all individuals of a common lineage within the same grid cell). The second format is a fasta file containing the entire genome of all individuals. Association of sequences to individuals, linkage units, genes and coded traits is defined in the fasta headers. Output is stored at the beginning and end of a simulation and at user-definable intervals. The output considers the state of all non-seed individuals at those times.
